# Distinct aneuploid karyotypes are universally selected for across cancers

**DOI:** 10.1101/2025.07.25.666795

**Authors:** Lucija Tomašić, Shane A. Fiorenza, Thomas W. van Ravesteyn, Geert J.P.L. Kops, Nenad Pavin

## Abstract

Aneuploid karyotype evolution leads to immense variability in cancer patient prognosis, so characterizing relevant trends has been an important objective (1–9). Recurrent chromosome gains and losses as well as oncogenic mutations have been associated with specific cancers (10–14), but a general trend in aneuploidy evolution has not yet been identified. Here we show a universal selection principle for karyotypes composed entirely of disomies and trisomies that is conserved across many different cancer types and even in yeast. This finding is surprising enough on its own, as cancers do not typically have generalized trends, but we also see that these karyotypes exhibit a 3 times lower mutation rate of the cancer suppressor gene TP53 compared to other aneuploid karyotypes. We find that, in general, karyotypes composed of combinations of any two chromosome copy numbers, termed *binary karyotypes*, make up over three-quarters of the Mitelman database, which we explain by a 15% increased fitness relative to other aneuploid karyotypes. In conclusion, our results reveal binary karyotypes as a distinct category of aneuploid karyotypes that are tolerated by p53 and universally selected for independent of encoded genes or any other chromosome-specific feature, shifting the current paradigm of aneuploid cell classification.

---

Aneuploidy, a deleterious condition marked by an abnormal number of chromosomes, leads to developmental disorders and is present in 90% of solid tumors as well as 50–70% of hematologic cancers (1–4). Dynamics of aneuploid karyotype evolution vary significantly between patients and have profound implications in cancer therapeutic responses and clinical outcomes (5–7). Due to this variability, a lot of effort has been put into characterizing aneuploidy and finding trends in the evolution of aneuploid karyotypes (6–9). For example, alterations in whole chromosome copy numbers are considered by tracking individual gains and losses of entire chromosomes relative to euploid. This approach has been used to identify recurrent gains of chromosomes 7, 8, 13, and 20 in colorectal carcinoma (10) and recurrent gains of chromosome 21 in acute lymphocytic leukemia (11). Despite these being some of the most prominent recurrent gains, they occur in only a small fraction of patients, with just 15% of acute lymphocytic leukemia cases exhibiting chromosome 21 gains (11). Therefore, while recurrent gains and losses are useful for describing individual cancers, they are limited in their ability to describe general trends in aneuploidy, as they are not highly conserved even within the same cancer type.

To potentially reveal more general trends in aneuploidy across different cancer types, both distance from euploidy, calculated from total chromosome number, and gene mutation are used (7,8,12–17). It was shown that karyotypes with 2N, 3N, or 4N total chromosome number dominate over aneuploid karyotypes, which was explained by aneuploid cells having lower fitness due to karyotypic imbalance. Despite this, aneuploid cells are widely present in cancers, and it has been suggested that this occurs due to mutations of specific genes (12,13). The most prominent example is mutation of the TP53 gene (14,18). It has been found that aneuploid karyotypes with ploidy lower than 2 have an association with TP53 mutation (19). More generally, this mutation has been observed in approximately 40% of patients (7), suggesting that TP53 mutations alone cannot fully explain aneuploidy in cancer. However, a link between gene mutations and broader trends in aneuploid karyotype evolution is missing.

In this paper, we find that the vast majority of human cancers exhibit a striking global trend in whole chromosome copy number alterations. By using a macro-karyotype approach in combination with mathematical modeling, we show that karyotypes composed of disomies and trisomies dominate over all other karyotypes across many cancers. This subset of karyotypes, which are composed of combinations of only two chromosome copy numbers, are referred to as binary. Our model predicts that binary karyotypes have ~15% increased fitness over other aneuploid karyotypes. We see that predominance of binary karyotypes also appears in both healthy human cells and yeast, suggesting it may be fundamental in nature and relevant from a broader evolutionary point of view. Remarkably, we find that the TP53 gene is mutated as much as 3 times less often in binary karyotypes than other aneuploid karyotypes. These findings reveal a universal selection principle for binary karyotypes, and this subset of aneuploid karyotypes behaves as an intermediate between euploid and highly aneuploid that can have fitness and TP53 gene mutation rates comparable to diploid cells.

## Binary karyotypes dominate over mixed

We first explore the collective behavior of different copy numbers in karyotypes by using cancer patient data from the Mitelman database, which includes both leukemia and solid tumors (21,22). We use a macro-karyotype approach, which focuses on chromosome copy numbers and is agnostic to individual chromosome identity (Fig. 1a). This approach lets us directly visualize the karyotype distribution and collective distance from euploidy in a two-dimensional plot (Fig. 1b, Box 1). This facilitates easier visualization of large datasets and quantitative comparison of different datasets than traditional karyotype heatmaps (Fig. 1c). Plotting the same leukemia data using our macro-karyotype approach reveals a striking regularity in karyotype distribution that is not obvious when a typical karyotype heatmap is used, though it can be recognized by regrouping based on copy number combinations (Figs. 1d,e). We see that the vast majority of cells occupy the edges of the macro-karyotype plot with a maximum at the right vertex. In particular, 44% of data is located along the hypotenuse, which corresponds to karyotypes composed entirely of disomies and trisomies (Fig. 1e). We also find that 32% of data is composed entirely of monosomies and disomies, located along the horizontal axis, and 1% of data is composed entirely of trisomies and tetrasomies, located along the vertical axis. Here, we see that these karyotypes, a subset which we will refer to as *binary* since they arise from combinations of two different copy numbers, make up 77% of the overall data (Fig. 1d). Karyotypes that are not binary, i.e., that arise from combinations of 3 or more different chromosome copy numbers, are referred to as *mixed* and make up the remaining 23% of data. These data show that binary karyotypes make up the vast majority of data despite each type representing just (2/5)^22^ = 0.0000002% of all possible karyotype combinations, indicating a strong selection pressure among aneuploid karyotypes. Furthermore, since this selection pressure does not depend on individual chromosome identity, these results suggest a universal mechanism on copy number state selection, regardless of encoded genes or any other chromosome-specific feature.

**Figure 1.**
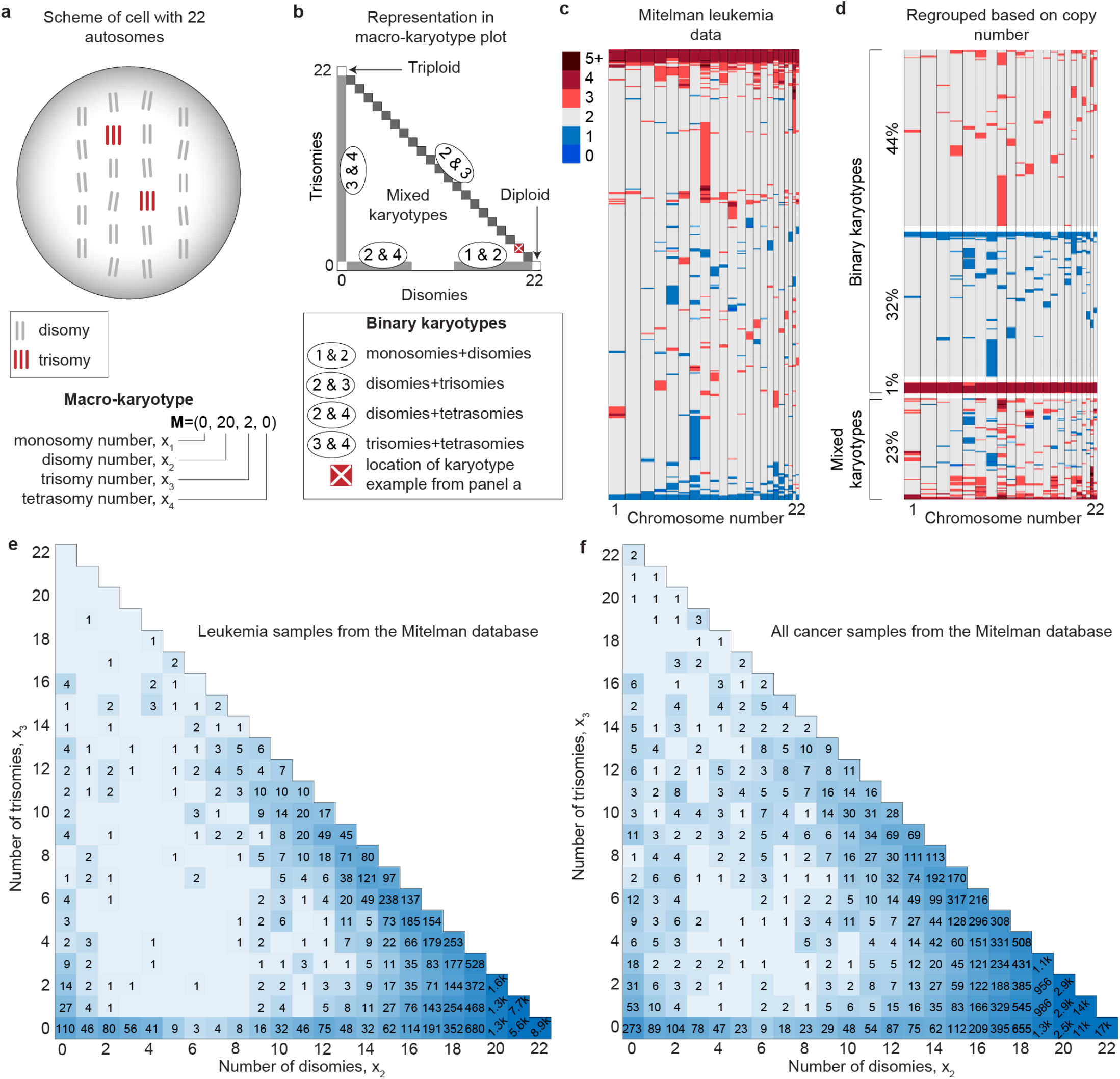
Macro-karyotype representation reveals regularity across many different cancer types. **a,** Scheme of cell with 20 disomies and 2 trisomies and its corresponding macro-karyotype. **b,** Graphical representation of aneuploid karyotypes obtained by plotting the number of disomies and trisomies on horizontal and vertical axes, respectively (triangle). This graphical representation formally corresponds to a projection of the macro-karyotype vector onto two-dimensional space, which we refer to as a macro-karyotype plot. Areas which correspond to four different types of binary karyotypes and the macro-karyotype example from (**a**) are highlighted (legend). **c**, Karyotype heatmap of 400 randomly-chosen leukemia patients from the Mitelman database, with each row representing a different patient. Copy number is indicated by color (legend). **d,** The same data from (**c**), but sorted into binary karyotype sub-groups to demonstrate their prevalence. **e** and **f**, Data of 33k patients with different types of leukemia (**e**) and 63k patients across all cancer types (**f**) from the Mitelman database shown in a macro-karyotype plot. The numbers in each square correspond to the amount of patient samples with the corresponding disomy and trisomy count. Shades of blue are given by a logarithmic function.

Given the stark preference for binary karyotypes in leukemia, we wondered if such a trend could be observed across different cancer types. We explore this by next looking at the entire Mitelman database, which includes aggregate tumor data from many different types of cancer, and we find that once again binary karyotypes are most common (Fig. 1f). Furthermore, if we run this analysis for each individual cancer type, we observe similar trends (Extended data Figs. 1, 2). Therefore, selection for binary karyotypes appears to be a universal trend across many cancer types.

### Binary karyotypes are more fit than mixed

To explore how binary karyotype distributions arise from an initially diploid state, we used mathematical modeling of macro-karyotype evolution to determine which mechanisms could result in the observed trends. We incorporate common mitotic errors into our model, such as single chromosome mis-segregation (23–26), whole genome duplication (17,27), and multipolar spindle cell division (28,29) (Fig. 2a, Methods). Single chromosome mis-segregations on their own can produce a variety of possible karyotypes, however, the number of binary karyotypes that arise from this mechanism alone is vanishingly small (Fig. 2b). Moreover, the addition of whole genome duplication results in a karyotype distribution that cannot be distinguished from those in Fig. 2b, indicating that another mechanism is needed. Based on these results, we propose that binary karyotypes have an increased fitness compared to mixed, similar to what has been established for euploid karyotypes. Implementing this idea allows our model to recreate the observed distributions for the hypotenuse and horizontal edge (Fig. 2c). Further, the addition of whole genome duplication now results in a karyotype distribution that occupies a distinct region of macro-karyotype space (Fig. 2d). Thus, our relatively simple hypothesis of binary karyotypes having increased fitness compared to mixed can explain the major experimental trends in both leukemia and aggregate cancer data (Figs. 1e, f). Using a quantitative comparison of the number of binary karyotypes versus mixed, our model estimates the fitness advantage of binary and euploid karyotypes to be 15% more than mixed on average (Fig. 2e).

**Figure 2.**
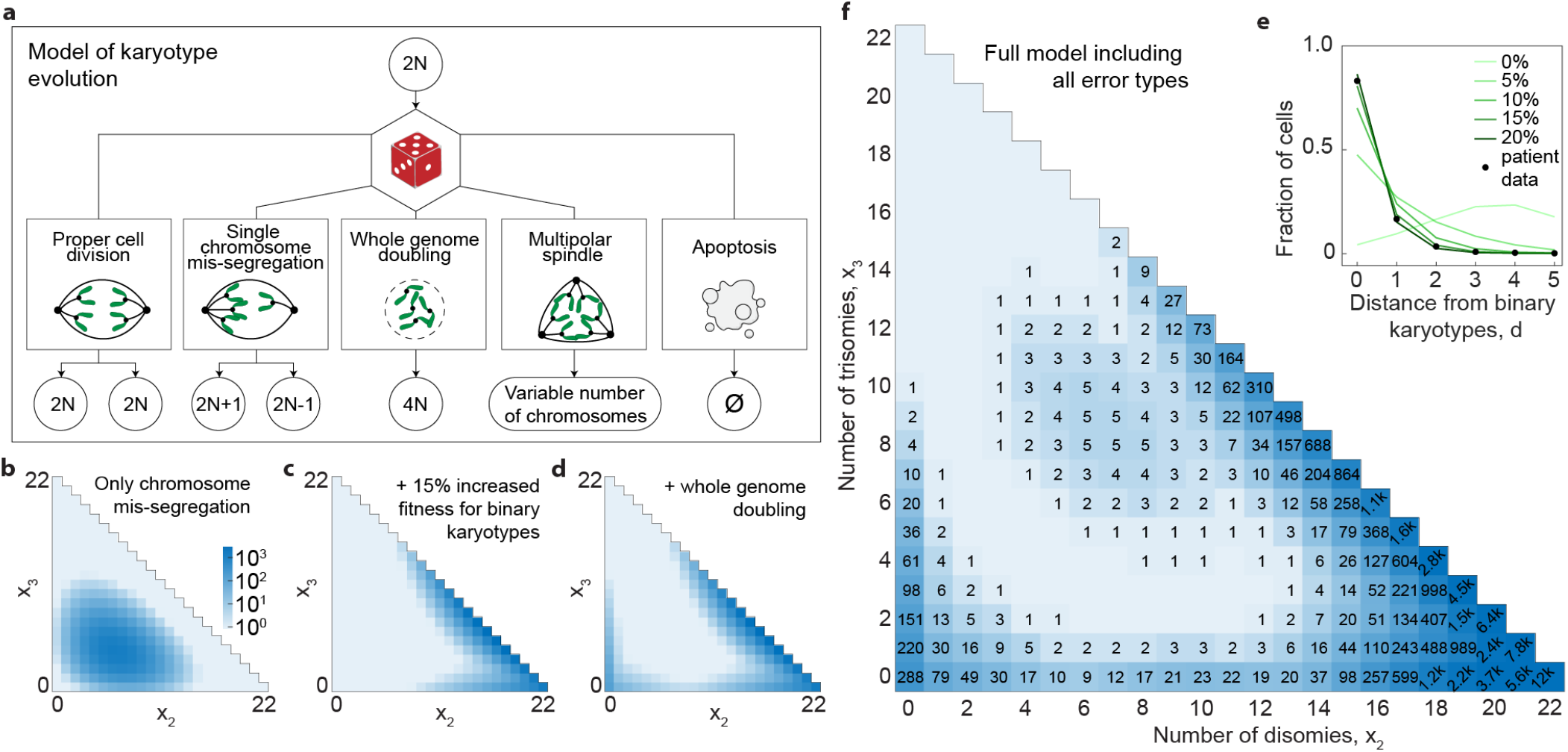
Theory shows each mitotic error type populates a distinct region in macro-karyotype space. **a**, Schematic of our mathematical model which depicts proper cell division (left rectangle), three common mitotic errors (middle rectangles), and apoptosis (right rectangle). Red dice indicates that cell fate is described by a stochastic process (Methods). **b**, Solution of the model with only single chromosome mis-segregations shown in a macro-karyotype plot. Parameter values are *p*_mis_ = 0.002 and *β* = 1 d^−1^, with the remaining set to zero. Legend indicates how shades of blue correspond to number of cells. **c**, Solution of the model with the addition of increased apoptosis of mixed karyotypes, 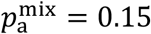, and slightly increased apoptosis of binary karyotypes composed of monosomies and disomies, 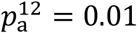. The remaining parameters are as in (**b**). **d**, Solution of the model when whole genome duplication is incorporated, *p*_WGD_ = 0.005, in addition to other processes described under (**c**). **e**, Fraction of cells at a given distance from binary karyotypes for solutions of the model with different apoptosis rates of mixed karyotypes (green lines) and patient data from Fig. 1**e** (black points). Legends shows 5 different values of parameter 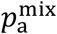. **f**, Solution of the model when all mitotic error types in (**a**) are included. Parameter values are the same as in plot (**d**), 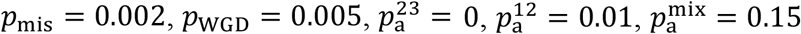, and *β* = 1 d^−1^ with the addition of *p*_MP_ = 0.002. Plot (**b**) shows distributions of karyotypes after 1000 days starting from initial diploid cells, whereas plots (**c, d**, and **f**) show the sum of 10 evenly-spaced timepoints over 100 days in addition to a single long-term datapoint after 500 days. Data is normalized at each timepoint so that the final aggregate plot has total sample number equal to those of the Mitelman database to allow for direct comparison.

Even though we were able to explain the experimental trends in leukemia without multipolar spindles, they are known to occur in other types of cancer (30,38). To investigate the role of multipolar spindles, we add to our model tripolar cell division that can occur following whole genome duplication. We see that multipolar cell division results in a broad region of mixed karyotypes that is distinct from other mechanisms (Fig. 2f). Therefore, multipolar cell division is a plausible explanation for the region of macro-karyotype space that appears when the entire Mitelman database, including solid tumors, is analyzed (Fig. 1f). Altogether, we see that these predominant mitotic errors are sufficient to recreate all trends observed in patient data, and each mitotic error populates a separate region in macro-karyotype space. Therefore, by leveraging the fact that each region in macro-karyotype space can be associated with a specific error, observed karyotype distributions in patient data can reveal the underlying mitotic error types.

### Cells evolve through binary karyotypes

We have shown that binary karyotypes are predominant in cancer samples (Figs. 1c, f), and our model explains this through them having increased fitness (Figs. 2c, d). To explore whether this trend is unique to cancers or more fundamental in nature, we next examined the karyotype evolution of healthy human cells and other organisms by reanalyzing published data using our macro-karyotype approach.

In the case of healthy human cells, which are typically diploid, aneuploidy was induced in cells and the karyotype distribution was tracked over time (31). In these experiments, aneuploidy is first induced in healthy breast cells, causing them to exhibit significant heterogeneity in karyotypes (Fig. 3a). In our macro-karyotype plot, this corresponds to a broad distribution of karyotypes. The karyotype distribution quickly reached a steady state after just 5-7 days, or two population doublings, and exhibited much less heterogeneity (Fig. 3b). Our macro-karyotype plots reveal that this steady state corresponds to karyotypes that are predominantly binary, either near tetraploid or diploid. The configuration of the steady state of karyotypes in these experiments, along with how quickly it was reached, is consistent with our hypothesis that there is a strong selection pressure for binary karyotypes. Thus, the similarity between experimental karyotype distribution and our model, which was used to explain patient cancer data, supports the idea that our hypothesis is relevant to healthy cells.

**Figure 3.**
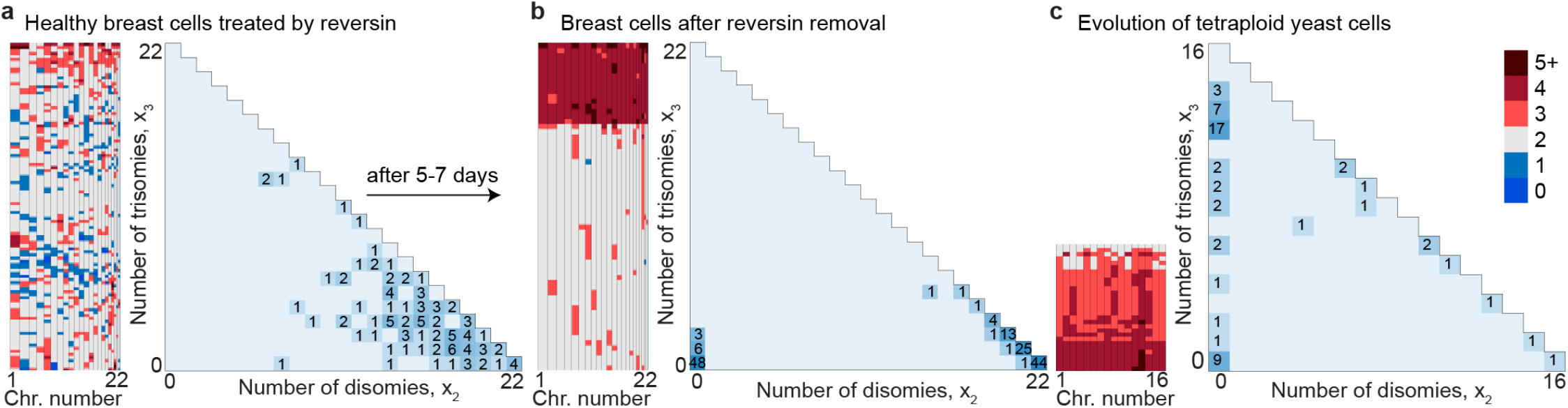
Healthy human cells and yeast evolve through binary karyotypes. **a,** (left) Karyotype heatmap of healthy breast cells following induced aneuploidy via reversin treatment from Ref. (31). (right) Data from left panel in a macro-karyotype plot, showing number of cells and corresponding shades of blue for a given number of disomies and trisomies. **b,** (left) Karyotype heatmap of healthy breast cells 5-7 days after reversin removal from Ref. (31), corresponding to approximately 2 population doublings. (right) Data from left panel in a macro-karyotype plot as in (**a**). **c,** (left) Karyotype heatmap of aggregate data of 35, 55, 250, and 500 generations after induced tetraploidization in yeast cells from Ref. (32). (right) Data from left panel in a macro-karyotype plot, showing that yeast cells evolve from tetraploid to diploid through binary karyotype intermediates.

To study binary karyotypes from a broader evolutionary perspective, we explore which general trends are conserved across the karyotype evolution of an organism that is distant from humans. We choose yeast as a model organism because it is a single-cell organism that has a substantially different spindle structure to humans. We analyze data from yeast karyotype evolution, which start from a tetraploid state and is tracked over many generations (32). Our macro-karyotype plot shows general trends in yeast evolution that exhibit a striking similarity to the karyotype evolution we observe in human cells. In particular, these data show that yeast evolve along the edges of the triangle, i.e., they predominately have a binary karyotype as they gradually transition from tetraploid to diploid (Fig. 3c). We see that in early generations, binary karyotypes with only trisomies and tetrasomies dominate. However, in later generations, binary karyotypes with only disomies and trisomies begin to appear and then increase over time. Altogether, these data show that binary karyotypes can facilitate evolution between two euploid states. Furthermore, the strong prevalence of binary karyotypes in a distant organism such as yeast supports the idea that our hypothesis is fundamental in nature.

### P53 tolerates binary karyotypes

So far we have identified trends in chromosome copy number alternates, but we also wanted to see if we can relate them to gene mutations. To explore this, we analyzed data from TCGA database since it includes information related to gene mutation (Fig. 4a). The most prominent candidate is TP53, which is the most frequently mutated gene in cancer because its inactivation facilitates tolerance of aneuploid karyotypes (18,33–35). To this end, we visualized the percent of samples in TCGA database with TP53 mutations in a macro-karyotype plot (Fig. 4b). We immediately notice that unaltered TP53 is more prevalent in binary and near-binary karyotypes, particularly those along the hypotenuse. In the center of the plot, corresponding to more mixed karyotypes, TP53 tends to be more frequently mutated. As expected, TP53 mutations are present across the entirety of macro-karyotype space, with the notable exception of karyotypes with only disomies and trisomies (Extended Data Fig. 3a,b). We also see that binary karyotypes with monosomies are more prone to increased rates of TP53 mutation as numbers of monosomies increases (Extended Data Fig. 3c), providing an explanation for why TP53 mutation is associated with lower ploidy (19). These results show that a clear link exists between binary karyotypes and TP53 mutation.

**Figure 4.**
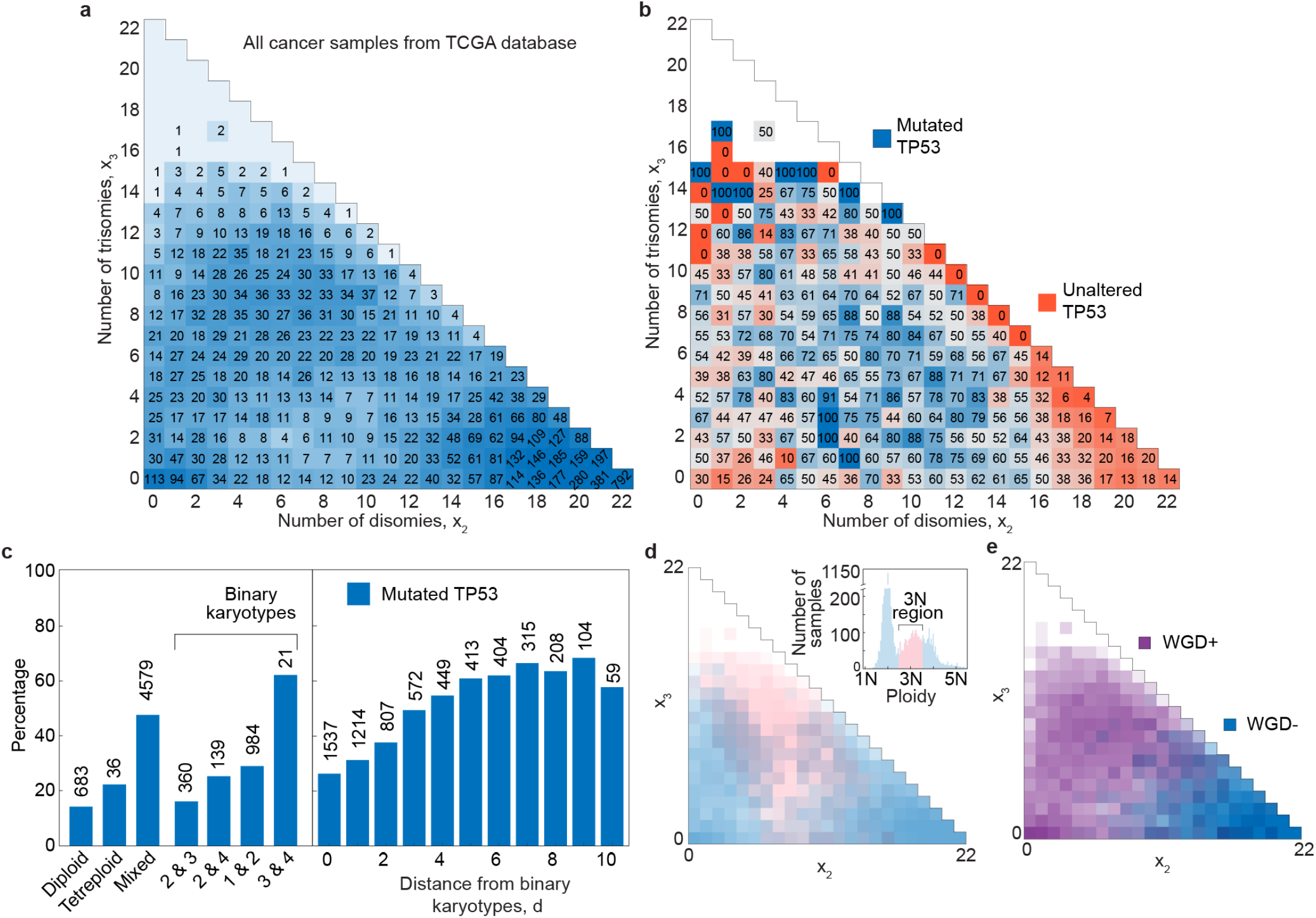
Binary karyotypes in TCGA database typically have unaltered TP53. **a,** All cancers from TCGA database shown in a macro-karyotype plot. **b,** Percent of samples from plot (**a**) with TP53 mutations. Low percentages (<50%) correspond to shades of orange, whereas high percentages (>50%) correspond to shades of blue. The boundary of these two regions (50%) appears in grey. **c**, (left) Bar plot showing percent of samples with TP53 mutations for diploid, tetraploid, and mixed karyotypes, as well as the four predominant types of binary karyotypes. (right) Percent of samples with TP53 mutation as a function of distance from binary karyotypes, with euploid karyotypes excluded. Numbers above bars denote total number of samples. This plot shows that the four predominant binary karyotypes in the left panel make up 98% of all binary karyotypes (*d* = 0 in right panel). **d,** Macro-karyotype plot and ploidy plot (inset) showing the same data from TCGA database as in panel (**a**), but with the data from 3N region, defined as being between 2.5N and 3.5N, highlighted in pink. **e,** The same plot as panel (**a**), but with karyotypes that have most likely undergone whole genome duplication (WGD+) highlighted in purple (Methods).

To gain a better understanding of the interplay between different subsets of karyotypes and TP53 mutation, we analyzed the percent mutation for the four predominant types of binary karyotypes in addition to diploid, tetraploid, and mixed karyotypes (Fig. 4c, left). We see that diploid karyotypes have the lowest rate of TP53 mutation as expected, whereas tetraploid and mixed karyotypes have rates that are 1.6 and 3.5 times larger as compared to diploid, respectively. Surprisingly, we see that aneuploid karyotypes composed of only disomies and trisomies have a rate of TP53 mutation that is lower than tetraploid and comparable to diploid. Karyotypes composed of only monosomies and disomies as well as disomies and tetrasomies have similar rates of TP53 mutation, approximately 2 times larger than diploid. The highest rate of TP53 mutation is observed for aneuploid karyotypes composed of only trisomies and tetrasomies, which we attribute to compounded errors associated with whole genome duplication or just small number statistics. These results show that binary karyotypes typically have substantially lower rates of TP53 mutation than mixed karyotypes, with those consisting of combinations of diploid and triploids being a pronounced example.

To further explore the difference between binary and mixed karyotypes, we plotted the TP53 mutation frequency as a function of distance from a binary state (Fig. 4c, right). We observe a consistent increase in mutations as distance from binary karyotype increases, i.e., as the karyotype becomes more mixed. Therefore, we propose that binary karyotypes are tolerated by p53, reinforcing our idea that they are distinct from other aneuploid karyotypes.

### Triploid karyotypes are underrepresented

It has been suggested that cancer cells frequently have near triploid karyotypes due to observed peaks at 3N chromosome number in ploidy plots for TCGA data (Fig. 4d inset, Refs. 16,17,36). Looking at TCGA data in macro-karyotype space, however, we see that no data is observed near the top vertex of the triangle, a region which corresponds to triploid and near triploid karyotypes (Fig. 4a). In fact, data only starts to appear for karyotypes that have up to 17 trisomies, indicating they are far from a triploid state which would consist of 22 trisomies in our analysis. By visualizing the exact distance from euploidy, our macro-karyotype approach reveals that cells predominately avoid the near triploid region.

We have shown that there is a negligible number of cells near the triploid region for TCGA database, raising questions regarding the nature of karyotypes that correspond to the 3N peak in ploidy plots. To answer this, we visualize the location of the 3N peak in macro-karyotype space by assigning a different color to karyotypes between 2.5N and 3.5N, corresponding to total chromosome numbers between 55 and 77 (Fig. 4d). This analysis reveals that the 3N peak in ploidy plots has no typical singular karyotype, but instead comes from more mixed karyotypes located across a broad region of macro-karyotype space. Furthermore, the vast majority of data that corresponds to the 3N peak comes from samples that have likely undergone whole genome duplication (Fig. 4e, Ref. 16).

Taken together, these results suggest that karyotypes which correspond to the 3N peak arise from a mechanism that involves genome doubling and produces a wide range of mixed karyotypes. A good candidate for this mechanism is multipolar cell division, which is known to occur more often in cells that have undergone whole genome duplication (38,39). We also observed in the exploration of our model that multipolar cell division following whole genome duplication results in a broad distribution of karyotypes similar to those that correspond to the 3N peak in ploidy plots (compare Figs. 2f and 4d). Therefore, our theory suggests that multipolar cell divisions are a likely candidate for causing the mixed karyotype distributions that appear as peaks near 3N in ploidy plots.

## Discussion

Here we have identified a subset of aneuploid karyotypes, referred to as binary karyotypes, that are universally selected for across different cancer and cell types (Figs. 1, 3). We have shown that cells can significantly deviate from a euploid state so long as they retain a binary karyotype, which our model predicts have 15% increased fitness relative to other aneuploid karyotypes (Fig. 2). This expands upon previous work that proposed euploid karyotypes are more fit due to gene dosage balance and that cell fitness depends on distance from a euploid state (1,15,37). However, here we have found that distance from euploidy is not the full story. This is illustrated by our observation that diploid states acquire either trisomies or monosomies as they evolve, whereas combinations of all three are observed less often even if they have the same distance from euploidy. This same prevalence of binary karyotypes is seen in healthy human cells and yeast, showing that it is relevant from a more general evolutionary perspective. Therefore, binary karyotypes reveal a hierarchy within aneuploidy that was previously hidden, introducing a fundamentally different perspective within aneuploid karyotype classification.

We have shown that binary karyotypes are characterized by low rates of TP53 mutation, which was as much as 3 times lower than other aneuploid karyotypes (Figs. 4b, c and Extended data Fig. 3). We have also found that as karyotypes deviate from a binary state, TP53 mutations become more prevalent (Fig. 4c). The TP53 gene is the most commonly mutated gene in cancer cells, because the protein it encodes promotes apoptosis of aneuploid cells (33,34). Therefore, the low rates of TP53 mutation in binary karyotypes may indicate that binary karyotypes are tolerated by the cell and function similarly to euploid cells, thereby circumventing the surveillance of p53. Based on this, we propose that binary karyotype may facilitate a balance between oncogenic and tumor-suppressor genes such that p53 does not need to be inactivated. We hypothesize that this is the driving factor of the relative fitness increase for binary karyotypes reported in this paper, opening an interesting idea for future studies.

Our results with TP53 show that a direct link between gene mutation and karyotype configuration is possible, introducing a new way of analyzing trends in cancers. The genes PIK3CA and KRAS have been identified as frequent alterations in cancer and are the most common after TP53 (7). Aneuploidy score and fraction of genome altered have been used to study how degree of aneuploidy is related to mutations in these genes (43,44). However, these techniques ignore the contribution of chromosome copy numbers and karyotype configuration as a whole. Reanalyzing mutations of these genes from a macro-karyotype perspective has the potential to reveal new trends and provide additional information related to their role in aneuploidy evolution of cancers.

In conclusion, we have shown that binary karyotypes are selected for across many different human cancer types. Even though this is surprising enough on its own, as cancers are not known to have many generalized trends, here we have also shown that binary karyotypes are a distinct subset with rates of TP53 mutation that are lower than other aneuploid karyotypes. We observed the same prevalence of binary karyotypes in yeast, an organism that is significantly distant from humans and has entirely different spindle, chromosome, and cell architecture. The striking similarity in observations between these two distant organisms suggests that binary karyotypes may be a fundamental occurrence in karyotype evolution that is independent of gene-or chromosome-specific features. By acting as an intermediate between highly aneuploid and euploid karyotypes, binary karyotypes may facilitate stable evolution between euploid states and even enable new branching points in evolutionary trajectories. Altogether, our results reveal a subset of aneuploid karyotypes that are universally selected for, challenging the dichotomy of simple aneuploid versus euploid categorization.

## Methods

### Macro-karyotype representation and description of binary karyotypes

In our theory, we consider karyotypes that are composed exclusively of whole chromosomes, where the *i*-th chromosome has *c_i_* copies which are strictly positive integers, i.e., *c_i_* ∈ ℕ, since we do not include nullisomies. For a cell with *N* different chromosomes, the entire karyotype is described by a N-dimensional vector **K** ≡ (*c*_1_, …, *c_N_*). To reduce the dimensionality of vector space, we instead use a macro-karyotype approach, **M**(**K**) ≡ (*x*_1_, …, *x_L_*) (20). Here, the vector component *x_j_* denotes the number of chromosomes,given by 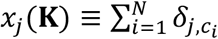, where *j* is the number of copies and *δ* is the Kronecker delta. The upper limit of the copy number is denoted by index *L*, which we chose to be as small as possible without compromising the accuracy of our approach. In this work, we use *L* = 5 in our model and *L* = 7 when analyzing experimental databases. The number of different chromosomes per cell is conserved, i.e., *x*_1_ + ⋯ + *x_L_* = *N*, as a consequence of bounded chromosome copy numbers, 1 ≤ *c_i_* ≤ *L*.

Binary karyotypes are a subset of aneuploid karyotypes which are composed entirely of combinations of two different chromosome copy numbers. In a macro-karyotype representation, a binary karyotype composed entirely of copy numbers *α* and *β* can be described as any vector with components obeying 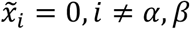, yielding 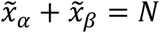. Along with binary karyotypes, we also define the distance from a binary state of *α* and *β* for any aneuploid karyotype through *d_αβ_* = *N* − (*x_α_* + *x_β_*). From a biological perspective, the minimum distance from any binary karyotype is most relevant, and therefore we use *d* = min(*d_αβ_*) for all combinations *α, β* ∈ [1, *L*] as a chosen measure.

### Mathematical model

To understand the potential physical mechanisms responsible for observed trends, we introduce a mathematical model of cell division and aneuploidy progression (Fig. 2), building upon previous work (36,37,45). We expand the model described in Ref. (20) using a set of coupled ordinary differential equations (ODEs) to model the karyotype progression of a population of cells. To calculate how the number of cells of a given macro-karyotype, *n*(**M**), changes in time, *t*, we use a mean-field approach. The dynamic equation that includes mis-segregations, whole genome doubling, multipolar spindles, and apoptosis is given as

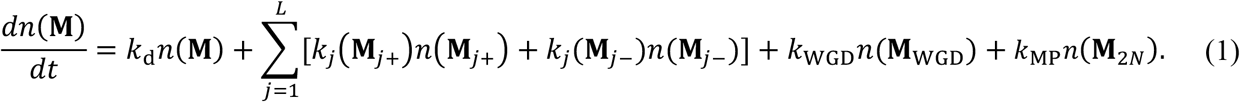

The first term in Eq. (1) represents events which do not change the karyotype of the mother cell, and the rate of this process is given as

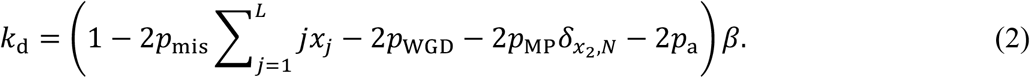

Here, the rate of cell division, *β*, defines average cell lifetime as ln 2 /*β*. After this amount of time, cell fate in our model is determined by a stochastic process where mis-segregation of a single chromosome, whole genome doubling, multipolar cell division, and apoptosis occur with probabilities *p*_mis_, *p*_WGD_, *p*_MP_, and *p*_a_, respectively. Note that Eqs. (1) and (2) are valid only for *p*_mis_ ≪ 1. For detailed calculations, see Ref. (20). The probability of apoptosis is smaller for binary karyotypes,

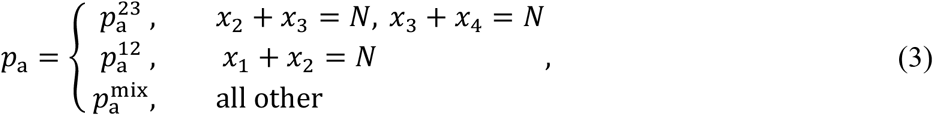

whereas all other probabilities are constants. The Kronecker delta term ensures that only tetraploid cells can undergo multipolar cell division. The second term in Eq. (1) describes whole chromosome mis-segregations, which occur with rate

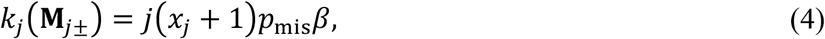

and change the macro-karyotype of daughter cells relative to the mother cell through the relation **M**_*j*±_ = **M** + **e***_j_* − **e**_*j*±1_. Here, the unit vector **e***_j_* has components with value 1 in the *j*th coordinate and 0’s elsewhere such that plus and minus signs denote relative gains and losses, respectively. Note that vectors **M**_*j*±_ with indices that exceed the bounds [1, L] or have any negative component, i.e., *x_j_* < 0, do not belong to macro-karyotype space and thus are not included in calculations. The third term in Eq. (1) describes whole genome doubling, which occurs with rate

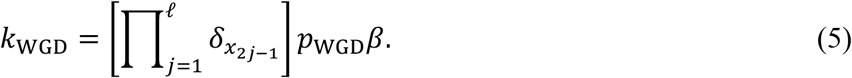

Here, the upper limit is always an integer calculated by using the floor function, *ℓ* = ⌊*I*/2⌋, and the Kronecker delta ensures that daughter cell have an even number of each chromosome upon whole genome doubling. The macro-karyotype of the mother cell is given by **M**_WGD_ = (*x*_2_, *x*_4_, …, *x*_2*P*_, 0, …, 0). The last term in Eq. (1) describes multipolar cell divisions, which in our model occur when a diploid cell, given by macro-karyotype **M**_2*N*_ = (0, *N*, 0, …,0), undergoes whole genome doubling that is immediately followed by formation of a multipolar spindle. Because tripolar spindles occur the most frequently (40), we only calculate chromosome distributions of daughter cells arising from this type of cell division. The rate of this event is calculated as:

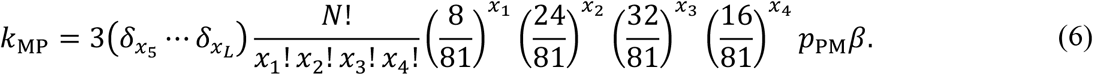

This rate comes from considering the combinatorics of chromosome arrangement prior to division into three daughter cells. Our model assumes that each daughter cell has zero nullisomies, i.e., gets at least 1 copy of each chromosome, so only combinations of 1 to 4 copies are considered. For this same reason, the Kronecker delta terms prevent combinations that would result in 5 or more copies. We also assume that each chromosome can establish bi-orientation with any two spindle poles out of the three with equal probability. Taken together, Eqs. (1)–(6) fully define the set of coupled ODEs that we solve numerically in order to explore evolution of karyotypes over time in our model.

### Data sources

We analyzed chromosome-level aneuploidy using publicly available datasets from the Mitelman and The Cancer Genome Atlas (TCGA) databases. Copy number data of individual TCGA samples were analyzed by ASCAT (41), obtained from the ASCAT Github repository (42). To ensure high-quality estimates of absolute copy number, we filtered out samples with estimated tumor purity that was either below 40% or exactly 100%. Samples with 100% purity typically represent normal tissue profiles, whereas low-purity samples often yield noisy or unreliable copy number calls. Copy number data of individual Mitelman samples were analyzed by CytoConverter (22), obtained from Mitelman database (21). Karyotype entries were included only if they provided complete, unambiguous whole-genome karyotype descriptions. Specifically, we filtered out: (1) karyotypes labeled as incomplete (those ending with “inc.” in the cytogenetic nomenclature), (2) karyotypes where the total chromosome number is given as a range rather than a specific value, and (3) samples flagged by CytoConverter with the warning that some chromosomes are unaccounted for.

In order to compare patient data with our mathematical modeling, where copy number is represented by an integer value, we rounded all data from databases. To this aim, for each autosome in a sample, chromosome copy number was calculated by weighing each segmental copy number value by its total segment length and then taking the statistical mode of all data. This approach ensures that the most representative copy number is assigned to each chromosome.

To show a link of macro-karyotype distribution with TP53 mutation and whole-genome doubling, additional analysis was performed. TP53 mutation status was downloaded from cBioPortal and associated with corresponding sample karyotype, whereas whole-genome doubling status for TCGA samples was taken directly from ASCAT output.

## Acknowledgements

We thank Kruno Vukušić and members of the N.P. and Iva Tolić groups for their constructive comments on the manuscript, as well as Ivana Šarić for assistance with assembling the figures. L.T. thanks Geoff Macintyre for useful discussion. This work was funded by the European Research Council (ERC-SyG 855158, N.P., and G.J.P.L.K.) and the European Union’s Horizon Europe research and innovation programme under the Marie Skłodowska-Curie Actions (MSCA 101151485, S.A.F.). Co-funding was provided by the Croatian Science Foundation (HRZZ IP-2019-04-5967, N.P.) and the Croatian Government and the European Union through the European Regional Development Fund—the QuantiXLie Center of Excellence (KK.01.1.1.01.0004, N.P.).

## Contributions

N.P. designed and supervised this study. L.T. and N.P. developed the theory. L.T. developed all code, performed research, and analyzed all data. S.A.F., T.V.R., and G.J.P.L.K. contributed with ideas and discussions. The manuscript was written by L.T., S.A.F., and N.P. with feedback from T.V.R. and G.J.P.L.K.

## Data availability

All relevant data supporting the findings of this study are provided within the article and its Extended Data files. Raw images used in this work are available from the corresponding authors upon reasonable request.

## Code availability

The code used to process patient samples from databases and for the numerical computations of the theoretical model are available upon request.

## Box 1.

**Visualizing aneuploidy in macro-karyotype space**

**Challenges associated with dimensionality of aneuploid karyotypes**

A major barrier to providing a comprehensive explanation of karyotype evolution in cancer is the sheer number of possible karyotype combinations. Human cancers, for example, have 22 autosomes with typically up to 5 copies each, and a simple calculation tells us this results in 5^22^ ≈ 2,400,000,000,000,000 combinations, making systematic study of all karyotypes impossible. Typical karyotype heatmaps display full karyotype information, but the resulting plots can be difficult to interpret visually due to the large dimensionality of the space, making direct comparison of different datasets challenging (Fig. 1d). This problem is exacerbated when using very large datasets, as it becomes infeasible for all information to be viewed at once. Therefore, multiple methods have been developed to represent different pertinent features in an easily-accessible manner, but they necessarily sacrifice some information in the process (16,32). For example, ploidy plots use total chromosome number to estimate distance from euploidy, but they lose information related to chromosome configuration. An approach that addresses this intractable dimensionality while still retaining key information of karyotype configuration is therefore necessary. An ideal candidate is a macro-karyotype approach, which reduces the dimensionality of karyotype space while retaining exact distance from euploidy (20).

**Representing macro-karyotype space in two-dimensional plots**

Rather than keeping track of each individual chromosome, a macro-karyotype approach focuses on chromosome copy number, which significantly reduces the dimensionality of the data (Fig. 1a, Methods). We can plot different chromosome copy numbers on each axis in a plot, allowing us to visualize the data. For optimal visualization, we use two chromosome copy numbers in order to produce a two-dimensional plot. In this work, we choose disomies and trisomies in particular, since they are predominant in the datasets we analyze. Alternatively, any combination of two chromosome copy numbers can be considered based on the needs of a given dataset, and sets of different combinations can be used to give a more comprehensive view of a single dataset.

To understand how to interpret macro-karyotype plots, we review some key features for our choice of disomy number and trisomy number as the horizontal and vertical axes, respectively. In this case, the right corner corresponds to a diploid karyotype, and the top corner corresponds to triploid (Fig. 1b). Other euploid karyotypes, such as haploid and tetraploid, appear at the origin in this representation. The hypotenuse, which connects the diploid and triploid corners, is composed entirely of combinations of disomies and trisomies and represents a region that is purely binary karyotypes. The other two edges include binary karyotypes but in theory do not exclusively correspond to them. However, in practice we see that the karyotypes in these regions are predominantly binary, with typical combinations labeled in Fig. 1b. Karyotypes that are mixed, i.e., those with 3 or more chromosome copy numbers, appear in remaining central region of the triangle. Typically, we observe that as distance from the edges increase, karyotypes become more mixed, allowing us to define a quantitative measure of distance from binary karyotypes (Methods).

## Extended Data

### Extended Data figures with legends

**Extended Data Figure 1.**
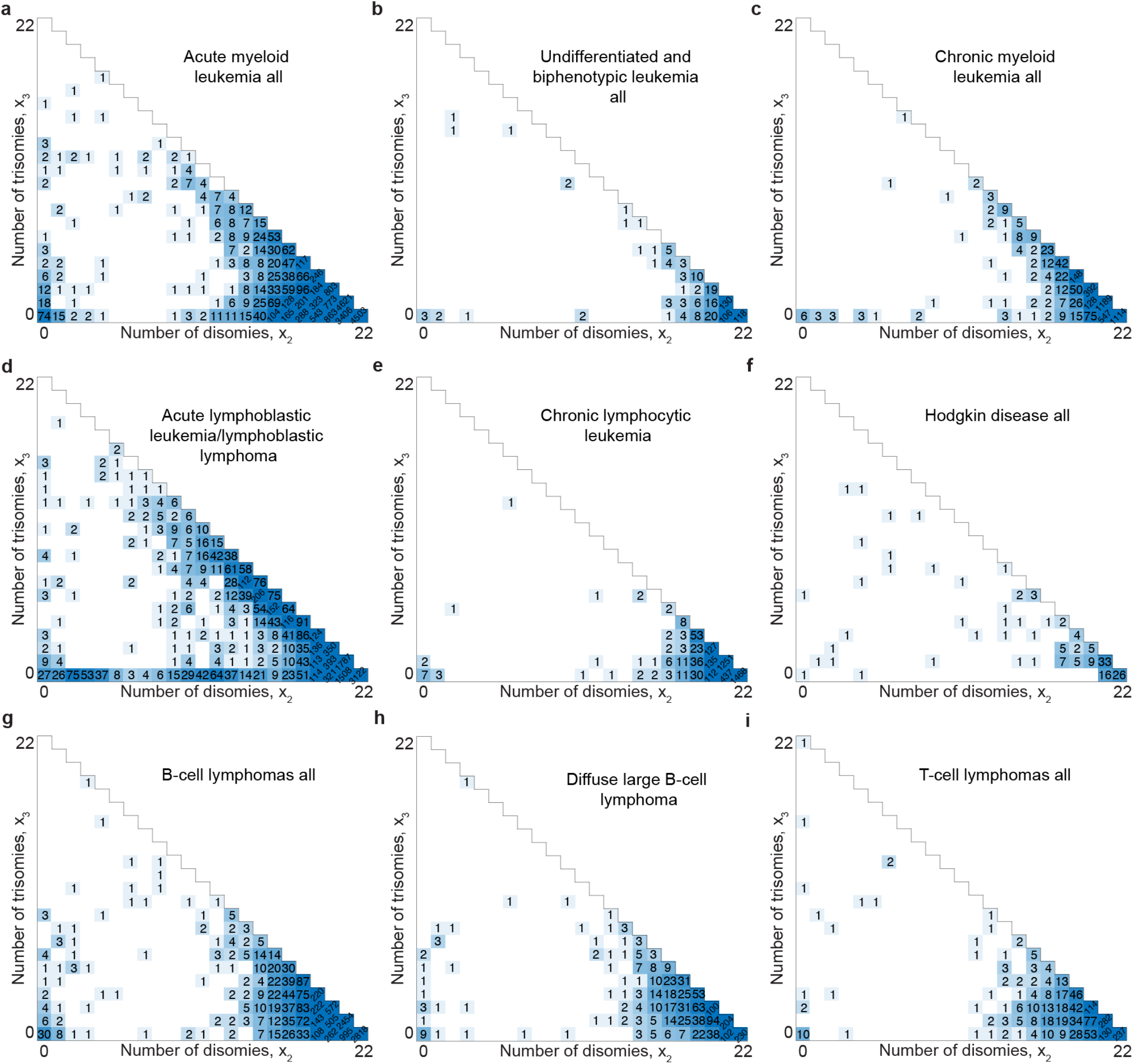
Macro-karyotype plots of patient samples for different leukemias and lymphomas from the Mitelman database. Macro-karyotype plots of patient-derived karyotypes from the Mitelman database illustrating chromosome copy number distribution across various hematological malignancies. **a**-**e**, Plots for the five most common leukemias. **f**-**i**, Plots for the four most common lymphomas. The numbers in each square correspond to the amount of patient samples with the corresponding disomy and trisomy count. Shades of blue are given by a logarithmic function. Cancers with a low number of datapoints, defined as having less than 10 aneuploid samples, are excluded.

**Extended Data Figure 2.**
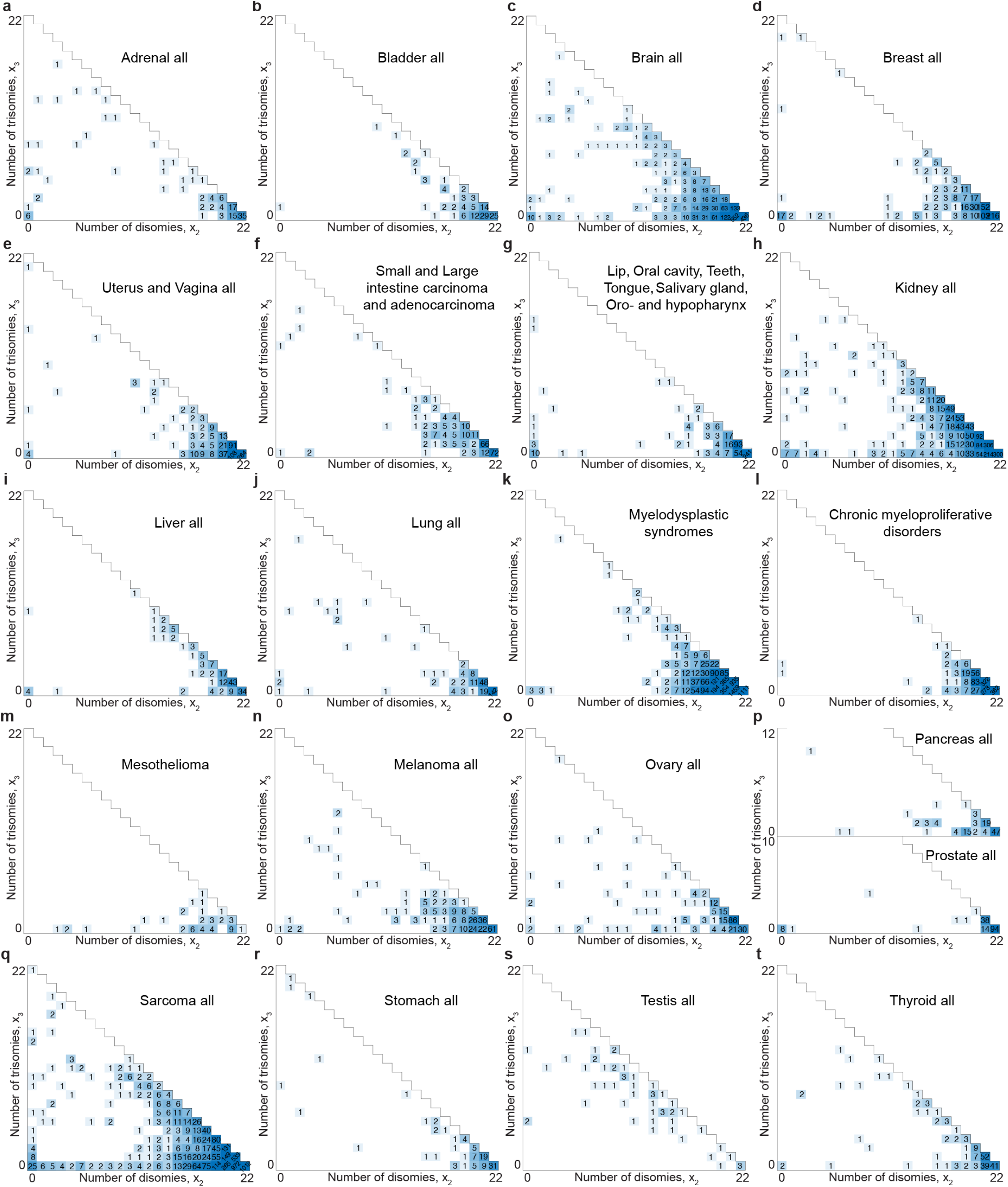
Macro-karyotype plots of patient samples for different cancers from the Mitelman database. Macro-karyotype plots of patient-derived karyotypes from the Mitelman database illustrating chromosome copy number distribution across various solid tumors. **a**-**t**, Plots for each different cancer type. The numbers in each square correspond to the amount of patient samples with the corresponding disomy and trisomy count. Shades of blue are given by a logarithmic function. Cancers with a low number of datapoints, defined as having less than 10 aneuploid samples, are excluded.

**Extended Data Figure 3.**
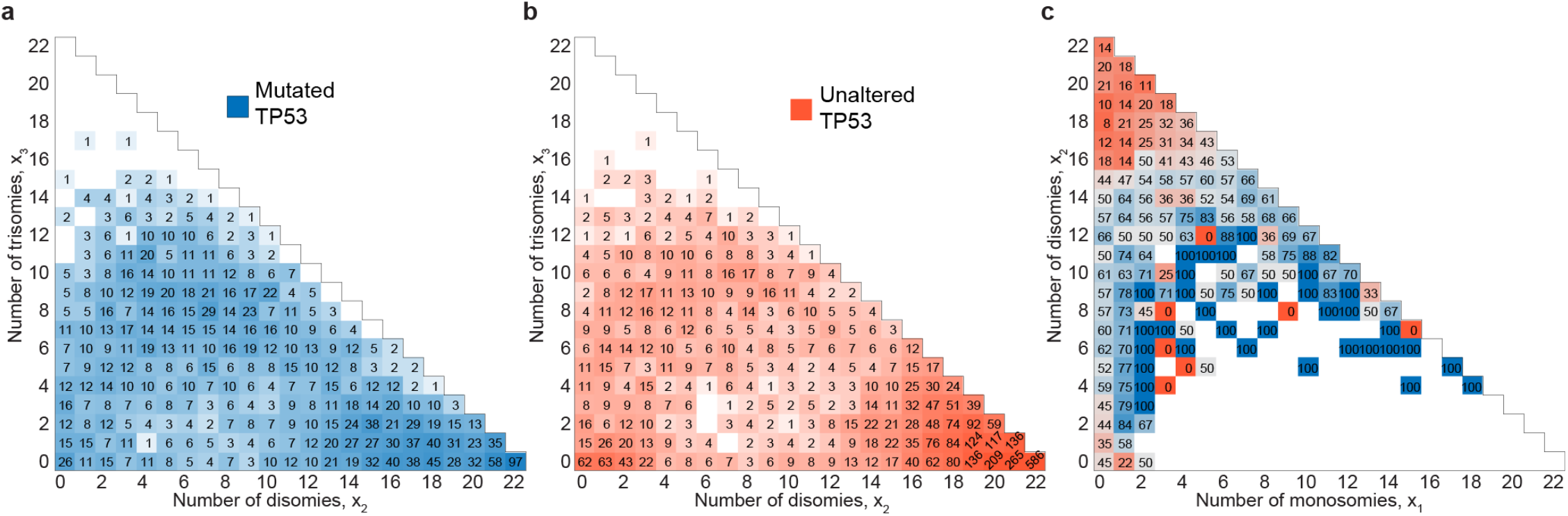
Macro-karyotype plots of TP53 mutations in patient samples from TCGA database. **a**, Number of samples from the plot in Fig. 4a that have TP53 mutations. Shades of blue are given by a logarithmic function. **b**, Number of samples from the plot in Fig. 4a without TP53 mutations. Shades of orange are given by a logarithmic function. **c**, Same data as in Fig. 4b, but represented in a plot with disomies and monosomies as the vertical and horizontal axes, respectively. In this plot, the hypotenuse corresponds exclusively to binary karyotypes composed of monosomies and disomies. Color coding is the same as Fig. 4b.

